# Repurposing the quinoline antibiotic nitroxoline to treat infections caused by the brain-eating amoeba *Balamuthia mandrillaris*

**DOI:** 10.1101/331785

**Authors:** Matthew T. Laurie, Corin V. White, Hanna Retallack, Wesley Wu, Matthew S. Moser, Judy Sakanari, Kenny Ang, Christopher Wilson, Michelle R. Arkin, Joseph L. DeRisi

**Affiliations:** Department of Biochemistry and Biophysics, University of California San Francisco, San Francisco, California, USA; California State University Monteray Bay, Seaside, California, USA; Department of Pharmaceutical Chemistry, University of California San Francisco, San Francisco, California, USA; Small Molecule Discovery Center, University of California San Francisco, San Francisco, California, USA; Chan Zuckerburg Biohub, San Francisco, California, USA

## Abstract

*Balamuthia mandrillaris* is a pathogenic free-living amoeba that causes a rare but almost always fatal infection of the central nervous system called granulomatous amoebic encephalitis (GAE). Two distinct forms of *B. mandrillaris* – a proliferative trophozoite form and a non-proliferative cyst form, which is highly resistant to harsh physical and chemical conditions – have been isolated from environmental samples worldwide and are both observed in infected tissue. Patients suffering from GAE are typically treated with aggressive and prolonged multi-drug regimens often including the antimicrobial agents miltefosine and pentamidine isethionate. However, survival rates remain low and studies evaluating the susceptibility of *B. mandrillaris* to these compounds and other potential therapeutics are limited. To address the need for more effective treatments, we screened 2,177 clinically-approved compounds for *in vitro* activity against *B. mandrillaris*. The quinoline antibiotic nitroxoline, which has safely been used in humans to treat urinary tract infections, was identified as a lead compound. We show that nitroxoline inhibits both trophozoites and cysts at low micromolar concentrations, which are within a physiologically relevant range. We compare the *in vitro* efficacy of nitroxoline to drugs currently used in the standard of care for GAE and find that nitroxoline is the most potent and selective inhibitor of *B. mandrillaris* tested. Furthermore, we demonstrate that nitroxoline prevents *B. mandrillaris-mediated* destruction of host cells in cultured fibroblast and primary brain explant models also at physiologically relevant concentrations. Together, our findings indicate that nitroxoline is a promising candidate for repurposing as a novel treatment of *B. mandrillaris* infections.

**Importance:** *Balamuthia mandrillaris* is responsible for hundreds of reported cases of amoebic encephalitis, the majority of which have been fatal. Despite being an exceptionally deadly pathogen, *B. mandrillaris* is understudied, leaving many open questions regarding epidemiology, diagnosis, and treatment. Due to the lack of effective drugs to fight *B. mandrillaris* infections, mortality rates remain high even for patients receiving intensive care. This study addresses the need for new anti-amoebic drugs using a high-throughput screening approach to identify novel *B. mandrillaris* inhibitors. The most promising candidate identified was the quinoline antibiotic nitroxoline, which has a long history of safe use in humans. We show that nitroxoline kills *B. mandrillaris* at physiologically relevant concentrations and exhibits greater potency and selectivity than drugs commonly used in the current standard of care. The findings we present demonstrate the potential of nitroxoline to be an important new tool in the treatment of life threatening *B. mandrillaris* infections.

## Introduction

The opportunistic protist pathogen, *Balamuthia mandrillaris* causes rare but life threatening infections of the central nervous system (CNS), termed *Balamuthia* or granulomatous amoebic encephalitis (GAE) (1, 2). Onset of the disease is gradual and chronically develops over a few weeks to months in both immunocompromised and immunocompetent individuals worldwide (1, 3). Presenting clinical symptoms include but are not limited to fever, vomiting, neck stiffness, headache, nausea, personality changes, and seizures (1, 3). These symptoms are nonspecific, overlapping with symptoms caused by more common brain infections such as bacterial and viral meningitis as well as non-infectious neuroinflammatory syndromes. Cutaneous presentation is less common and can produce symptoms ranging from painless swelling to ulceration and formation of large lesions (4–6). Infections involving other organs ranging from the lungs to the eye have also been documented (7, 8). Thus, *B. mandrillaris-induced* encephalitis often goes unrecognized and diagnosis is frequently only made postmortem. Several hundred cases have been reported, however, the actual disease burden is likely underestimated (2, 9).

While systematic ecological studies of *B. mandrillaris* have not been performed, free-living *B. mandrillaris* have been isolated from water, soil, and dust across all continents (10–18). Cases of human and animal infections are also reported on all continents but are most common in South America and the southern United States (reviewed in Bravo and Seas, 2012; H Diaz, 2011; and Centers for Disease Control (CDC) 2008) (19–35). *B. mandrillaris* is thought to be transmitted by inhalation of contaminated aerosols or exposure via broken skin (36). Fatal amoebic encephalitis has also occurred after solid organ transplantation (37, 38). Pathogenesis is believed to involve hematogenous spread to the CNS through penetration of the blood-brain barrier and amoebae are frequently observed around blood vessels (3, 36).

Free-living amoebae such as *Sappinia diploidea, Acanthamoeba spp*. and *Naegleria fowleri* can also cause infection of the CNS with very poor prognosis (2, 39). *Acanthamoeba* and *Balamuthia* are the most similar and are classified in the same eukaryote super group *Amoebozoa: Acanthamoebidae* (40). The mode of infection employed by *Sappinia diploidea* and *Acanthamoeba spp*. is thought to be similar to *Balamuthia*, as encephalitis caused by these genera progresses over several weeks to months and is associated with water or soil contact. On the other hand, acute encephalitis caused by *N. fowleri* is distinct and specifically associated with recreation in warm freshwater environments with presumed neuroinvasion of the amoeba by passing up the nose through the cribiform plate to the brain. All of these pathogenic amoebae have a proliferative trophozoite form and a dormant, thick-walled cyst form; the cyst form is notoriously more resistant to antimicrobials (41–46) and a variety of abiotic stressors such as ultraviolet light (47–49). Such attributes, along with the fact that drug sensitivities differ among genera, species, and strains of free-living amoebae have complicated studies in drug discovery (50).

Infection of the CNS by *B. mandrillaris* is almost always fatal and no specific and highly successful treatment regimen is known (44, 51). The CDC recommends the following drugs for treatment of *B. mandrillaris* CNS infection: pentamidine isethionate, miltefosine, fluconazole, flucytosine, sulfadiazine, azithromycin and/or clarithromycin (52). *In vitro* studies with the CDC-recommended drugs show little to no inhibition of amoebic growth by fluconazole, sulfadiazine and flucytosine while azithromycin, pentamidine isethionate, miltefosine, and voriconazole (fluconazole derivative) exhibit amoebicidal or amoebistatic activity (11, 41, 50). Current treatments for *B. mandrillaris* CNS infections employing experimental combinations of these drugs have produced inconsistent outcomes including survival in some cases and fatality in others (7, 19, 23, 27, 29, 30, 34, 53–61). As the efficacy and specificity of current treatments remains uncertain, there is a clear need to identify additional drugs that can improve patient outcomes.

The goal of this study was to identify, from a set of clinically-approved compounds, candidates that have the potential to be repurposed for treatment of *B. mandrillaris* infections. Here, we screened a library of 2,177 clinically-approved compounds and found that the quinoline antibiotic nitroxoline (8-hydroxy-5-nitroquinoline) exhibits amoebicidal activity at low micromolar concentrations, well within the range of estimated plasma concentrations achieved with recommended oral dosing (62–64). Through direct *in vitro* comparisons, we find that nitroxoline is a substantially more potent and selective inhibitor of *B. mandrillaris* than three commonly used GAE treatments currently recommended by the CDC. In addition to killing *B. mandrillaris* trophozoites, nitroxoline also causes encystment and substantially delays recrudescence of active amoebae, which is significant considering the rapid decompensation of patients suffering from GAE. Nitroxoline, with its ease of delivery and favorable pharmacodynamic properties, has the potential to be an effective treatment for GAE in singularity or in combination with drugs in the current standard of care.

## Methods

### Human Cell Lines

Hep-G2 (ATCC HB-8065), U87 (gift of Jonathan Weissman lab), H4 (ATCC HTB-148), and HEK-293T (ATCC CRL-3216) cells were cultured in Dulbecco’s Modified Eagle’s Medium (Gibco) containing 10% (vol/vol) FBS (Gibco), 2 mM L-glutamine, 100 U/mL penicillin/streptomycin (Gibco), and 10 mM Hepes buffer. HFF-1 cells (ATCC SCRC-1041) were cultured in DMEM containing 15% (vol/vol) FBS, 2 mM L-glutamine, 100 U/mL penicillin/streptomycin, and 10 mM Hepes buffer (Gibco). All mammalian cell incubation steps were carried out at 37°C with 5% CO_2_. All cell lines were tested for mycoplasma contamination using the Lonza Mycoalert mycoplasma detection kit (Lonza).

### *Balamuthia mandrillaris* propagation, handling and encystment

*Balamuthia mandrillaris* (ATCC PRA-291) were maintained axenically in 150 cm^2^ flasks (Corning) containing modified Cerva’s medium (axenic medium) with the following formulation: 20 g of Bacto Casitone (Difco), 68 mL of 10x Hank’s balanced salt solution (Gibco), 10% fetal bovine serum, and 1x penicillin-streptomycin (200 UI/mL – 200 μg/mL) (65). Axenic growth of *B. mandrillaris* resulted in 2-3×10^5^ amoebae/mL in log phase. Following findings of a previous study, encystment of *B. mandrillaris* was induced by galactose exposure (66). *B. mandrillaris* trophozoites were grown to log phase in axenic media and galactose was added to a final concentration of 12% (v/v). Amoebae were cultured in the induction medium until trophozoites were no longer observed (approximately 10 days). Galactose-induced cysts transition back to trophozoites after approximately three days of incubation in galactose-free axenic medium. Therefore, all assays with cysts were completed in the induction medium. To quantify amoebae for use in experiments, actively growing trophozoites or recently induced cysts were centrifuged at 3,000 rpm for 5 minutes, resuspended in axenic medium and counted with a disposable hemocytometer (SKC, Inc.). All incubation steps for *B. mandrillaris* growth were carried out at 37°C with 5% CO_2_.

### Primary drug screening of *Balamuthia mandrillaris* trophozoites *in vitro*

Screening of a clinically-approved library of compounds, compiled by the Small Molecule Discovery Center (SMDC) at the University of California San Francisco was completed at 20 μM in 0.2% DMSO. All 2,177 drugs in this library were stored as 10 mM stocks dissolved in 100% DMSO (Sigma-Aldrich) at −20°C. *B. mandrillaris* amoebae were resuspended in axenic medium and distributed into opaque 384-well plates (Corning) at a density of 3,000 amoebae per well using a BioMek NX liquid handler. Negative control wells were treated with vehicle only (0.2% DMSO in axenic medium) and positive control wells simulating total destruction of amoebae were seeded with amoeba lysate generated by 3 consecutive freeze-thaw cycles. Following 72 hours of incubation, 30 μL of CellTiter Glo reagent (CTG; Promega) was added to each well. The luminescence was measured with a Promega GloMax-Multi+ plate reader at 2, 4, and 8 hours after CTG addition. The percent inhibition of *B. mandrillaris* was calculated based on the CTG luminescence measurements of treated wells relative to both positive and negative controls using the following equation: % inhibition = 100 - 100*((test well intensity - positive control intensity)/(negative control - positive control)). The B-Score, a plate-based statistical approach for correcting row, column, and edge effects, was also calculated for each compound in the library (67). Raw luminescence measurements and computed inhibition values are displayed in Table S1A. Hits were determined to be compounds with a B-score of approximately 5 or above and percent inhibition of approximately 40% or above (Table S1B). Hit compounds that are approved only for topical administration or veterinary use were not tested in secondary screening (highlighted in yellow in Table S1B and Fig 1B). Compounds identified in the primary screen that did not exhibit activity upon repurchase of fresh compound were removed from consideration in the screening funnel.

**Figure 1:**
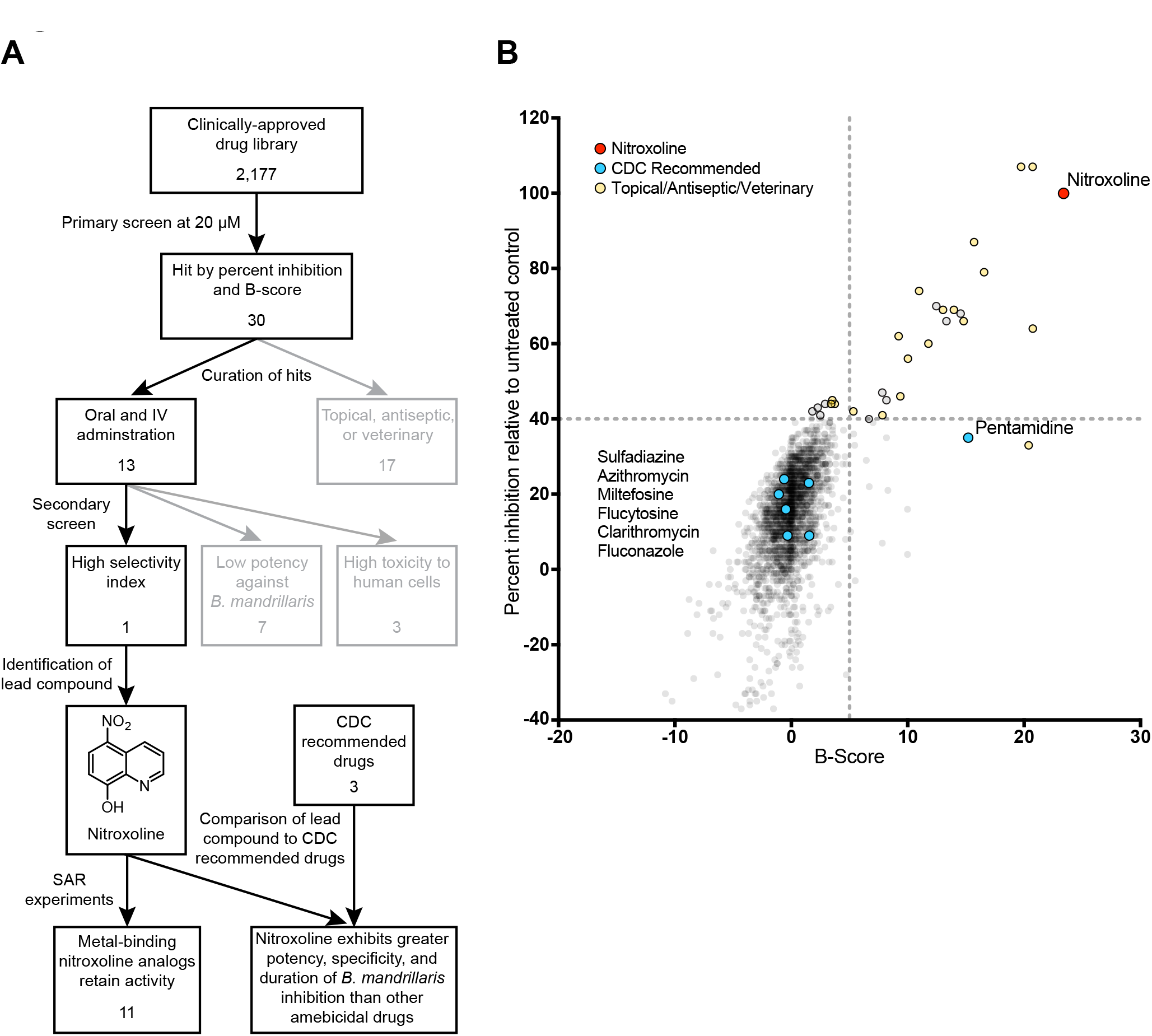
(A) Workflow for high-throughput screening of clinically-approved compounds for in vitro activity against *B. mandrillaris*. A primary screen of 2,177 clinically-approved compounds yielded 30 hits meeting the percent inhibition and B-score criteria, of which only 13 candidates were available for oral or IV administration (Fig. 1B). Secondary screening identified only one novel lead compound, nitroxoline, which displayed high selectivity for inhibition of *B. mandrillaris* viability (Table S1C). Structure-activity relationship (SAR) experiments show that 11 of 12 nitroxoline analogs tested with potential metal binding domains remain active against *B. mandrillaris*, suggesting that metal binding plays a role in the mechanism of inhibition by nitroxoline (Fig. 2). Comparison of nitroxoline to three drugs recommended by the CDC for treatment of *B. mandrillaris* CNS infections (pentamidine isethionate, miltefosine, and azithromycin) indicates that nitroxoline is the most potent and specific inhibitor of *B. mandrillaris* of the compounds tested (Fig. 3, 4 and Table 1). (B) Plot of percent inhibition relative to untreated controls and B-score measured for each compound in a library of 2,177 clinically-approved compounds. Raw data used to calculate these values is compiled in Table S1A. Drugs recommended by the CDC for treatment of GAE are highlighted in blue. Drugs that are classified as antiseptic, topical, and/or have not been used in humans are shown in yellow. The quinoline antibiotic nitroxoline, which was the top hit identified in this screen, is highlighted in red.

### Secondary drug screening against *B. mandrillaris* trophozoites, HFF-1 and H4 human cells *in vitro*

Secondary screening of hit compounds consisted of dose-response experiments to measure the toxicity of each compound to *B. mandrillaris* trophozoites, HFF-1 human fibroblast cells, and H4 human neuroglioma cells. Four 384-well plates were prepared prior to drug addition: one opaque plate (Corning) seeded with 3,000 HFF-1 cells per well in 60 μL complete media, one opaque plate seeded with 2,000 H4 cells per well in 60 μL complete media, one opaque plate containing 30 μL axenic media, and one clear plate (Corning) containing 30 μL axenic media. Opaque plates were used to measure the viability of trophozoites using the CTG assay and clear plates were used for microscopic examination of cyst formation. HFF-1 and H4 plates were seeded 24 hours prior to drug addition. The selected primary screen hits were added from 10 mM DMSO stocks into wells of all four test plates to reach final concentrations ranging from 0.06 μM to 30 μM (10 concentrations, 2-fold serial dilution). Negative control wells received concentrations of DMSO corresponding to the amount of DMSO in each tested drug well (reaching a maximum of 0.3% DMSO). *B. mandrillaris* trophozoites were resuspended in axenic media and added to the plates containing drug dilutions in axenic media at a density of 3,000 amoebae per well in a final volume of 60 μL media. All plates were incubated for 72 hours. Throughout the incubation period, the clear-bottom *B. mandrillaris* plate was monitored for large-scale changes to population encystment in response to drug treatment. After the 72 hour incubation, 30 μL of CTG reagent was added to all wells of the opaque assay plates and luminescence was measured with a Promega GloMax-Multi+ plate reader at 2, 4, and 8 hours after CTG addition. IC_50_ values for inhibition of *B. mandrillaris* viability were determined using GraphPad Prism 4-parametric sigmoidal curve fitting model, with bottom and top constraints set to 0 and 1 respectively (Table S1C).

### Nitroxoline structure-activity-relationship experiments

Nineteen commercially available analogs of nitroxoline were selected for structure-activity-relationship (SAR) experiments based on variance in functional groups that may play a role in the observed mechanism of action (compound sourcing and data in Table S2). *B. mandrillaris* were grown to log phase axenically and plated at 4,000 amoebae per well in opaque 96-well plates. Each nitroxoline analog was dissolved in 100% DMSO at 10 mM. Analog stocks were serially diluted in water to generate 8-point dilution series, which were then used to treat assay wells containing *B. mandrillaris* trophozoites at final concentrations ranging from 0.14 μM to 300 μM (8 concentrations, 3-fold serial dilution) in a final volume of 100 μL. After incubation for 72 hours, 50 μL of CTG was added to all assay wells. Luminescence was measured using the Promega GloMax Multi+ luminometer 2, 4, and 8 hours after CTG addition. IC_50_ values were determined using GraphPad Prism 4-parametric sigmoidal curve fitting model with bottom and top constrains of 0 and 1, respectively.

### Dose-response experiments with HFF-1, H4, U87, Hep-G2, HEK 293T cells and *Balamuthia mandrillaris* trophozoites and cysts

*B. mandrillaris* trophozoites were seeded at 4,000 amoebae per well into opaque and clear bottom 96-well plates (Corning). Homogenous populations of *B. mandrillaris* cysts generated by galactose induction were seeded at 4,000 amoebae per well into opaque 96-well plates. HFF-1 (fibroblast), H4 (glial), U87 (glial), HEK-293T (kidney), and Hep-G2 (liver) were seeded at 3,000 cells per well in opaque 96-well plates 24 hours prior to addition of drug. Stocks of nitroxoline (Selleck Chemicals) were dissolved in 100% DMSO at 10 mM. Stocks of azithromycin (Selleck Chemicals), pentamidine isethionate (Selleck Chemicals), and miltefosine (Selleck Chemicals) were dissolved in water at 10 mM. Drug stocks were serially diluted in water to generate 12-point dilution series, which were then used to treat assay wells containing *B. mandrillaris* or human cells at final concentrations ranging from 0.39 μM to 400 μM in 100 μL total well volume. Control wells were treated with vehicle (DMSO or water) at concentrations corresponding to the final vehicle concentrations in each drug dilution series. After incubation for 72 hours, 50 μL of CTG was added to all assay wells. Luminescence was measured with a Promega GloMax-Multi+ plate reader 2, 4, and 8 hours after CTG addition. All dose-response experiments were performed with at least three independent biological replicates. IC_50_ values were determined as previously described. To quantify encystment in treated and untreated conditions, amoebae in individual assay wells were resuspended thoroughly and the number of cysts and trophozoites in 10 μL samples were counted by hemocytometer (SKC, Inc.). Cysts and trophozoites are morphologically distinct, making them simple to distinguish with brightfield microscopy (example images shown in Fig. S1). Encystment assays were performed with three independent biological replicates.

### *Balamuthia mandrillaris* recrudescence assays

Populations of *B. mandrillaris* were diluted to 2.5×10^5^ amoebae in 10 mL media and were treated with 3.5, 7, 14, 28, 56, 84, and 112 μM nitroxoline, pentamidine isethionate or miltefosine and incubated for 72 hours. Following incubation, all remaining amoebae in each population (various mixtures of cysts and trophozoites) were pelleted at 3,000 rpm for 5 minutes and resuspended in drug-free HFF-1 medium. Each resuspended *B. mandrillaris* population was placed on a monolayer of HFF-1 cells that were seeded at 10^6^ cells per flask in 10 mL 24 hours prior to inoculation. At this cell density, untreated amoebae typically consume 100% of HFF-1 monolayers within 24 hours. Co-culture flasks containing amoebae and HFF-1 cells were incubated until 100% of the HFF-1 monolayer was consumed as observed by daily microscopic inspection or until the pre-determined endpoint of the experiment at 28 days post-*B. mandrillaris* inoculation. The day on which complete clearance of HFF-1 monolayer occurred was recorded for all conditions. All recrudescence assays were performed with three independent biological replicates. These methods were adapted from a minimum trophozoite amoebicidal concentration (MTAC) assay that was conducted with monolayers of MA104 monkey kidney cells (41).

### Primary brain tissue model

De-identified tissue samples were collected with previous patient consent in strict observance of the legal and institutional ethical regulations. Protocols were approved by the Human Gamete, Embryo, and Stem Cell Research Committee (institutional review board) at the University of California, San Francisco. Primary brain tissue samples were sectioned perpendicular to the ventricle to obtain slices 300 μm thick and ∼2.5 mm^2^ in surface area, using a Leica VT1200S vibrating blade microtome in artificial cerebrospinal fluid containing 125 mM NaCl, 2.5 mM KCl, 1 mM MgCl_2_, 1 mM CaCl_2_, 1.25 mM NaH_2_PO_4_. Explants were transferred to slice culture inserts (Millicell) in 6-well culture plates and cultured with media containing 66% Eagle’s Basal Medium, 25% Hanks balanced salt solution, 5% fetal bovine serum, 1% N-2 supplement, 1x penicillin-streptomycin, and glutamine, in a 37°C incubator with 5% CO_2_. 12 hours after plating, slices were inoculated with 10^4^ *B. mandrillaris* trophozoites in 20 μL of complete media containing vehicle (DMSO) or 35 μM nitroxoline, added dropwise to the slice surface. Media below the cell culture insert was adjusted to matching vehicle or nitroxoline concentrations. At 20 hours post-inoculation, media below the cell culture insert was replaced with fresh media containing no drug. At this time, amoeba on top of the insert surrounding and within the tissue were undisturbed. Brightfield and phase contrast images were captured during the live culture experiment at 4X, 10X, and 20X using an Evos FL Cell Imaging System. At 4 days post-inoculation, slices were gently fixed in 3.7% PFA for 4 hours at 4°C, then rinsed with PBS and stained for 2 hours at room temperature with DAPI (0.3 μM) in PBS with 1% Triton X-100, and then mounted with ProLong Gold Antifade Mountant (Thermo). Images of stained tissue were obtained with a Nikon Ti spinning disk confocal microscope at 20X magnification. Confocal z-stacks were projected and adjusted in ImageJ. Brightfield images were stitched using photomerge in Photoshop (Adobe). Brightfield time-lapse images were processed as movies in ImageJ.

## Results

### Identification of nitroxoline as an inhibitor of *B. mandrillaris* trophozoites *in vitro*

Due to the extremely high mortality rate associated with *Balamuthia mandrillaris* infections and the limited efficacy of current treatments, there is a clear need to identify additional therapeutic strategies to improve patient outcomes in these rare, but deadly infections. Here, we established replicating axenic cultures of *B. mandrillaris* (ATCC PRA-291) and screened 2,177 clinically-approved compounds for reduction of trophozoite viability following 72 hours of treatment at 20 μM. Compounds with high percent inhibition (∼40% or above) and B-score (∼5 or above) were nominated for secondary screening (Fig. 1A, Table S1A). Remaining compounds were annotated by class and delivery method to eliminate drugs only approved for topical delivery or veterinary use, which have low therapeutic potential (Table S1B). The selected candidate drugs were tested in dose-response assays with *B. mandrillaris* trophozoites as well as HFF-1 and H4 human cell cultures to confirm their activity and evaluate toxicity to human cells (Table S1C). Half-maximal inhibitory concentration (IC_50_) values indicated that only two compounds, pentamidine isethionate and nitroxoline, demonstrated adequate potency against *B. mandrillaris* without high toxicity to human cells. Since the amoebistatic activity of pentamidine has previously been described (45), we focused on further characterizing the novel activity of nitroxoline as the lead compound identified by this screen.

### Nitroxoline structure-activity-relationship experiments

Nitroxoline is currently in clinical use as an antimicrobial drug in certain European and Asian countries. The hypothesized mechanism of action (MoA) is as a metal chelator that disrupts biofilm formation (68). Nitroxoline and other 8-hydroxyquinolines have also previously demonstrated *in vitro* anticancer activity and free-radical metabolites are thought to be involved in cell death (69). To test which of these mechanisms is applicable in killing *B. mandrillaris*, we tested 19 commercially available analogs of nitroxoline for amoebicidal activity. While nitroxoline itself remained the most potent compound tested, there was a clear necessity for the 8-position hydroxyl group on the quinoline ring to retain high activity (Fig. 2, Table S2). As demonstrated by the activity of 8-hydroxyquinoline [**2**], the 5-position nitro group is not necessary for activity, while in contrast, retaining the nitro group without the hydroxy group [3] significantly decreases activity. Replacing the nitro group at the 5-position of the 8-hydroxyquinoline core with a variety of other functional groups, along with dual 5,7-position functionalization (compounds **4-13**) results in several active but less potent analogs with no clear trend due to aromatic electronic affects. The sulfonic acid functionalized variant [**13**] was the sole inactive compound among compounds **4-13**, which we speculate may be due to differences in uptake or permeability. Lastly, phenanthroline [**14**], a structurally similar bidentate metal binding ligand, also demonstrated activity against *B. mandrillaris*. Taken together, these results indicate a likely metal binding mechanism for the original nitroxoline compound.

**Figure 2:**
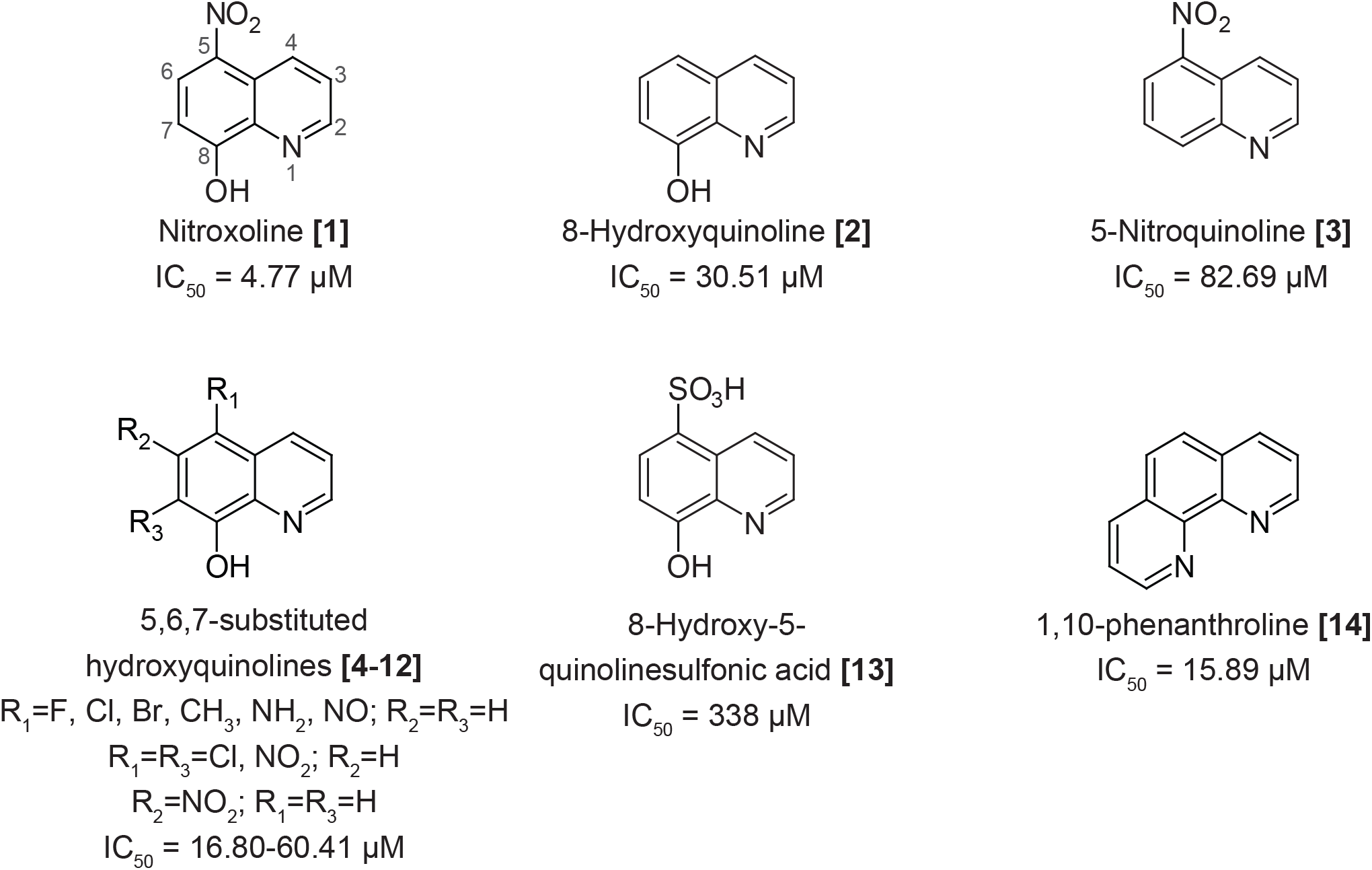
Structure-activity-relationship experiments suggest that nitroxoline inhibits *B. mandrillaris* through a mechanism related to metal binding. Structures and IC_50_ values are shown for nitroxoline and select analogs; additional compounds are shown in Table S2. Nitroxoline is made up of a quinoline core with a nitro group at the 5-position and hydroxyl group at the 8-position. Analogs lacking the 8-position hydroxyl group were generally inactive with IC_50_ values greater than 80 μM (e.g. 5-Nitroquinoline **[3]**). Twelve nitroxoline analogs with predicted metal binding activity were tested and of these, 1,10-phenanthroline **[14]** and 10 out of 11 compounds with an 8-position hydroxyl group (e.g. 8-Hydroxyquinoline **[2]**) were active with IC_50_ values ranging from 17-60 μM. The only inactive analog with an 8-position hydroxyl group was 8-Hydroxy-5-quinolinesulfonic acid **[13]**. Variance of the 5-position nitro group reduced potency compared to nitroxoline, but no trend related to aromatic electronic effects is apparent.

### Direct comparison of nitroxoline to standard-of-care drugs for GAE treatment

To evaluate nitroxoline as a potential drug to treat *B. mandrillaris* infections, we compared the *in vitro* performance of nitroxoline to pentamidine isethionate, miltefosine, and azithromycin, three drugs recommended by the CDC and commonly used in treatment of GAE (52). We performed side-by-side dose-response experiments to measure the efficacy of each drug against *B. mandrillaris* trophozoites and the toxicity to different human cell types using the following cell lines: HFF-1 (fibroblast), H4 (glial), U87 (glial), HEK-293T (kidney), and Hep-G2 (liver). Nitroxoline was the most potent inhibitor of *B. mandrillaris* trophozoites with an IC_50_ of 2.84 μM, compared to IC_50_ values of 9.14 μM, 63.23 μM, and 244.10 μM for pentamidine, miltefosine, and azithromycin, respectively (Fig. 3A). Nitroxoline was also the only drug with an IC_50_ against *B. mandrillaris* that was lower than the half-maximal cytotoxic concentration (CC_50_) for all cell lines tested. The average Log_10_ selectivity index (CC_50_ for human cell toxicity/IC_50_ for *B. mandrillaris* inhibition) of nitroxoline across all cell lines was 0.832, compared to 0.049, −0.102, and −0.409 for pentamidine, miltefosine, and azithromycin, respectively (Fig. 3B).

**Figure 3:**
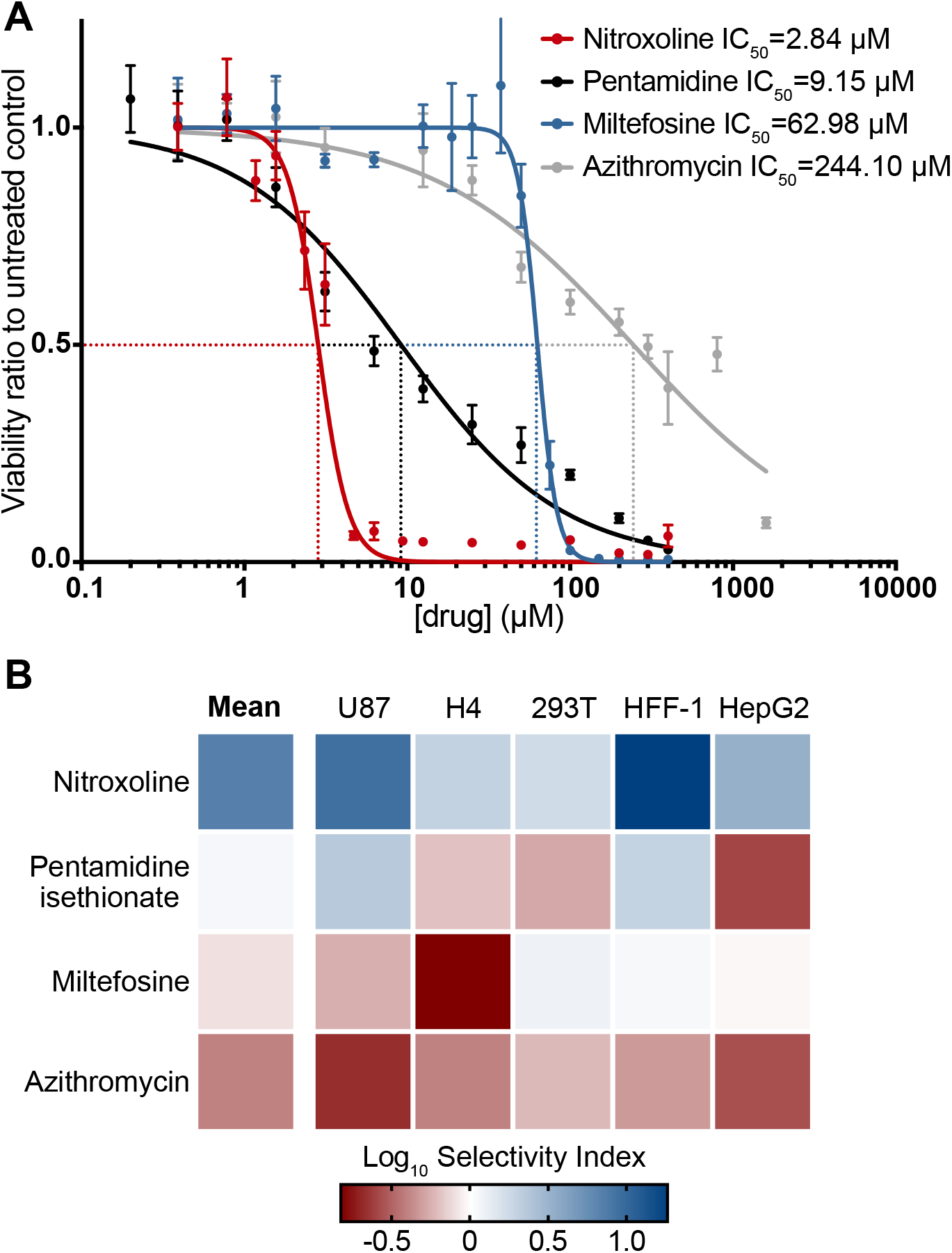
Potency and selectivity for inhibition of *B. mandrillaris* viability by nitroxoline, pentamidine isethionate, miltefosine, and azithromycin. (A) Dose-response curves show the effect of nitroxoline (red), pentamidine isethionate (black), miltefosine (blue), and azithromycin (grey) on the viability of *B. mandrillaris* trophozoite populations following 72 hours of treatment. Data points are means and standard errors of at least three independent biological replicates. Nitroxoline is the most potent inhibitor of *B. mandrillaris* viability with an IC_50_ of 2.84 μM. (B) Heat map showing the Log_10_ selectivity index (human cell CC_50_/*B. mandrillaris* IC_50_) for nitroxoline, pentamidine, isethionate, miltefosine, and azithromycin calculated from the ratio of human cell CC_50_ to *B. mandrillaris* IC_50_. Nitroxoline exhibited the greatest mean Log_10_ selectivity index at 0.832 and was the only drug with a positive Log_10_ selectivity index comparing *B. mandrillaris* inhibition to all cell lines tested.

### Encystment response of *B. mandrillaris*

In addition to dose-dependent reduction in *B. mandrillaris* trophozoite viability, we also observed a general increase in the ratio of cysts to trophozoites correlated with increasing concentration of some drugs. We investigated the propensity of nitroxoline, pentamidine isethionate, miltefosine, and azithromycin to induce encystment of *B. mandrillaris* by counting the number of cysts and trophozoites in culture samples following 72 hours of treatment with different drug concentrations. We observed a dose-dependent reduction in the number of trophozoites in the population for all four drugs. Nitroxoline and pentamidine isethionate caused an increase in both the total number and the proportion of cysts in the population, while no substantial number of cysts was observed at any concentration of miltefosine or azithromycin (Fig. 4A-D). Because encystment appears to occur as a response to certain drug treatments, we also assessed the ability of each drug to inhibit the viability of pre-formed *B. mandrillaris* cysts. We induced encystment by sustained exposure to 12% galactose and conducted dose-response viability measurements for each drug. Nitroxoline was again the most potent inhibitor of cysts with an IC_50_ of 15.48 μM compared to IC_50_ values of 26.26 μM, 76.48 μM, and 788.4 μM for pentamidine, miltefosine, and azithromycin, respectively (Fig. 4E). While nitroxoline, pentamidine, and azithromycin were considerably less potent inhibitors of cyst viability compared to trophozoites (Fig. 3A), miltefosine inhibited both forms of *B. mandrillaris* at similar concentrations.

**Figure 4:**
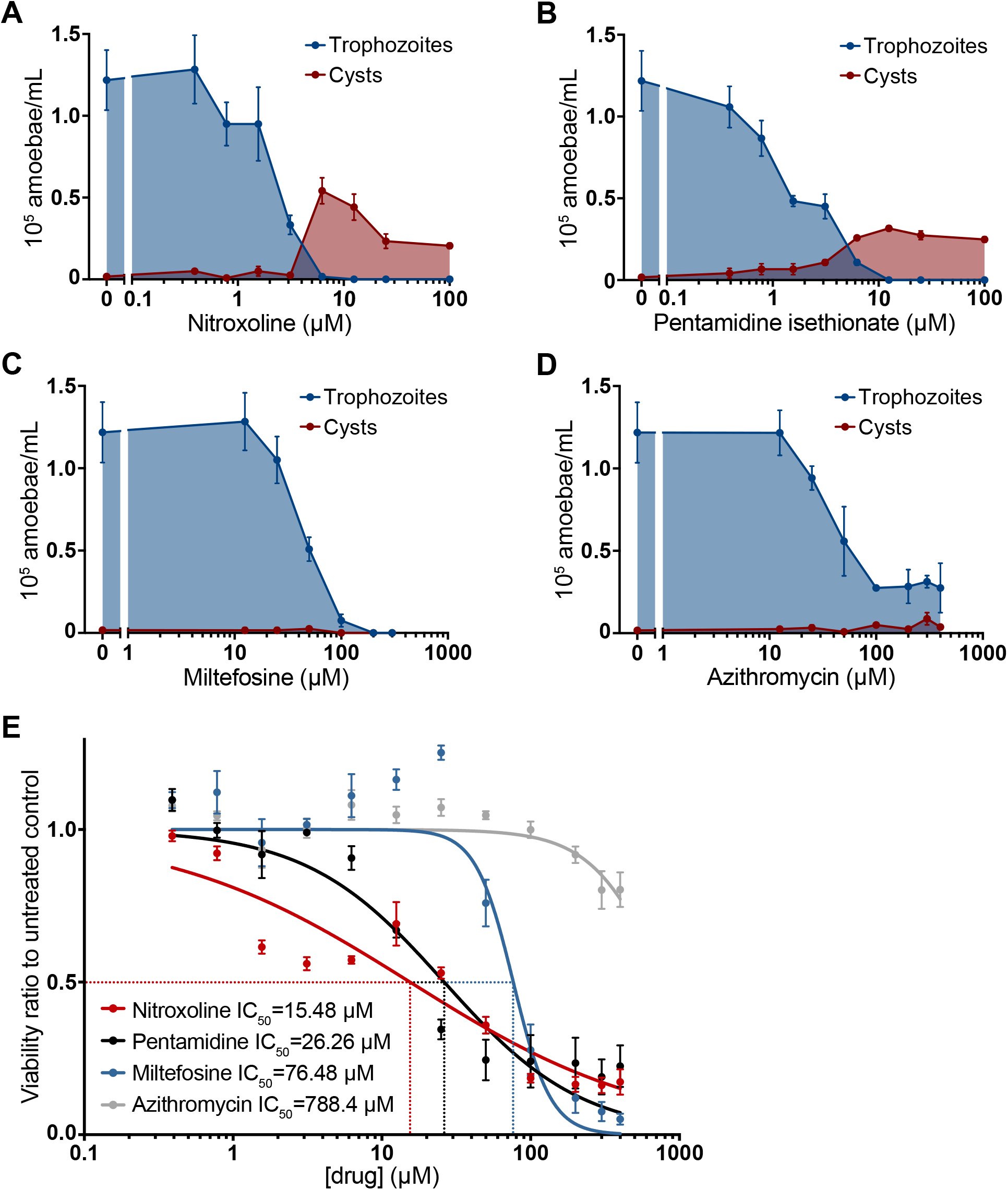
Relationship of drug treatment to *B. mandrillaris* encystment. (A-D) Changes in the number of trophozoites (blue) and cysts (red) in *B. mandrillaris* populations following 72 hours of treatment with various concentrations of nitroxoline (A), pentamidine isethionate (B), miltefosine (C), and azithromycin (D). Low micromolar doses of nitroxoline and pentamidine isethionate cause an increase in the total number of cysts observed in *B. mandrillaris* populations and an increase in the ratio of cysts to trophozoites. No increase in encystment is observed in *B. mandrillaris* populations treated with miltefosine or azithromycin. (E) Dose-response curve showing the effect of nitroxoline (red), pentamidine isethionate (black), miltefosine (blue), and azithromycin (grey) on the viability of pre-formed *B. mandrillaris* cysts. Nitroxoline is the most potent inhibitor of cysts with an IC_50_ of 15.48 μM. Compared to trophozoites (Fig. 2A), cysts are substantially less sensitive to all drugs except for miltefosine, which had a similar IC_50_ for inhibition of both *B. mandrillaris* forms.

### Delayed recrudescence of *B. mandrillaris* treated with nitroxoline

Nitroxoline and the tested standard-of-care drugs induce various combinations of distinct and intermediate *B. mandrillaris* phenotypes including death and encystment. Though it is assumed that drug-induced phenotypes such as encystment affect the rate of amoeba population growth and host cell destruction, the magnitude and duration of these effects are unknown. To assess how rapidly amoeba populations recover and proliferate following drug treatment, we developed a recrudescence assay wherein we treated *B. mandrillaris* trophozoite cultures with varying concentrations of nitroxoline, pentamidine, or miltefosine for 72 hours, removed drug, and then added the remaining amoebae to a monolayer of HFF-1 cells in the absence of drug. The post-treatment recovery time was measured as the number of days required for each *B. mandrillaris* population to clear 100% of the host cells. Treatments that completely eliminated *B. mandrillaris* populations were determined by observing no live trophozoites or destruction of host cells at any point during the 28 day experiment. We found that 7 μM and 14 μM pentamidine delayed recovery of *B. mandrillaris* by 1-2 weeks, but increasing the dose from 14 μM to 56 μM only delayed recovery by an additional 3 days. Consistent with the steep Hill slope observed in dose-response experiments, miltefosine caused very little delay to clearance time at 56 μM but completely eliminated *B. mandrillaris* populations at 84 μM. In contrast, nitroxoline delayed amoeba recovery by 2-3 weeks at low micromolar concentrations and completely eliminated *B. mandrillaris* populations at 28 μM.

### Protective effect of nitroxoline in a primary human brain tissue model

Findings from the recrudescence assays suggest that nitroxoline treatment may significantly impede destruction of host cells in the context of *B. mandrillaris* infection. To further explore this possibility, we performed experiments modeling *B. mandrillaris* infection and nitroxoline treatment in primary human brain tissue. Human cortical tissue slices were exposed to *B. mandrillaris* trophozoites and simultaneously treated with nitroxoline or vehicle (DMSO) for 20 hours before media was changed to remove drug or vehicle. Tissues were cultured for 4 days and then evaluated for damage by microscopic examination (Fig. 5 and S2). Untreated tissues show widespread damage following *B. mandrillaris* exposure including loss of distinct tissue edges and reduction of cell density (Fig. 5B). Large numbers of highly motile *B. mandrillaris* trophozoites can be seen at the edges of tissues and intermixed with human cells (Movie S1). In contrast, nitroxoline treated tissues did not show signs of *B. mandrillaris-mediated* destruction and appeared similar to uninfected tissues (Fig. 5A and C). Cysts with little to no motility are observed outside the boundaries of tissues (Movie S2). While these findings are qualitative, the large-scale differences in tissue morphology observed in this experiment are consistent with the possibility that nitroxoline has a protective effect towards host tissue in the context of *B. mandrillaris* infection.

**Figure 5:**
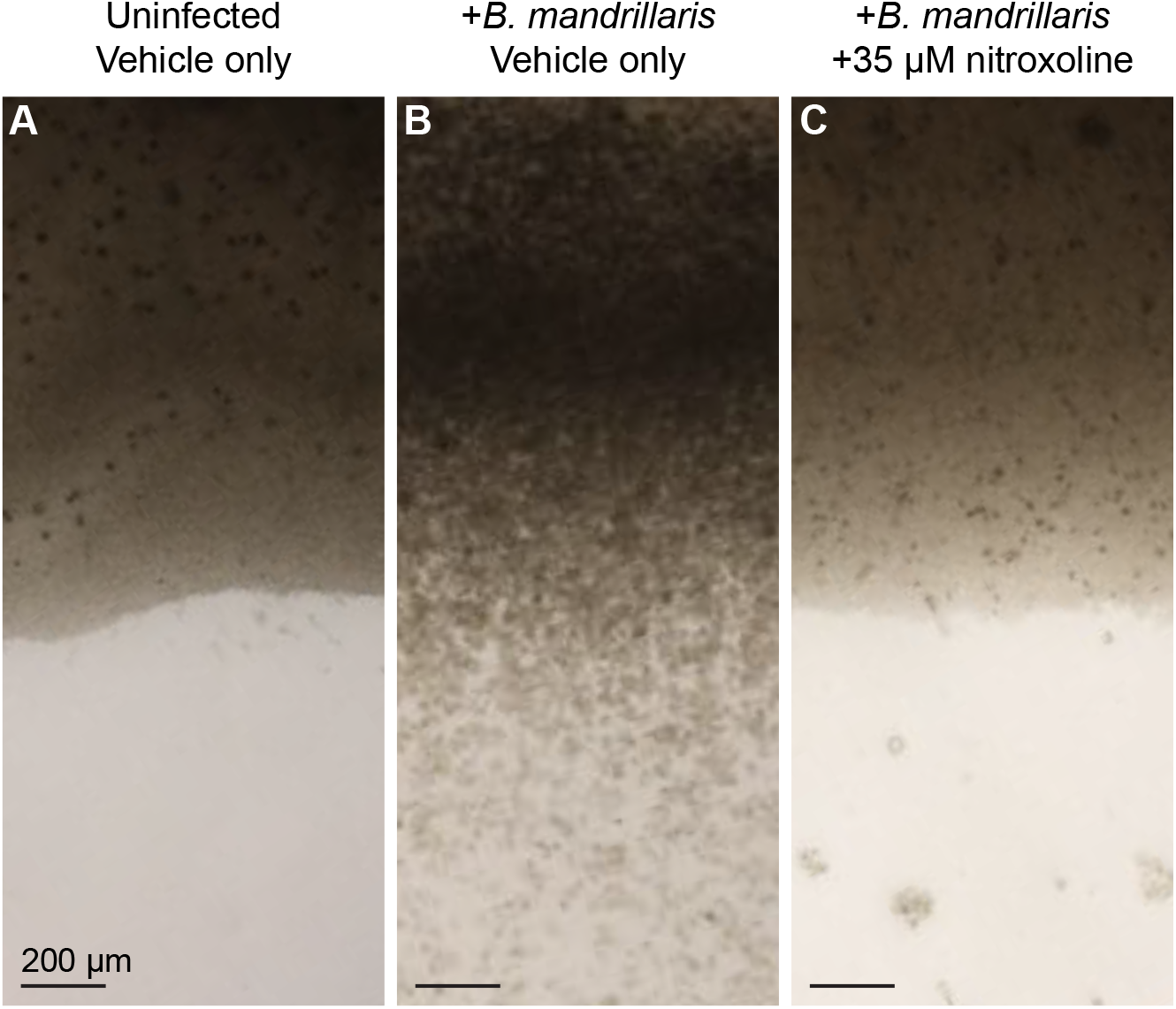
Nitroxoline prevents *B. mandrillaris-mediated* destruction of human brain tissue explants. Each panel shows an image representative of two tissues 96 hours after exposure to the indicated conditions. Media was changed at 20 hours post-infection to remove nitroxoline or vehicle. (A) Uninfected, untreated (vehicle only) tissues have distinct edges and maintain cell density throughout culture. (B) *B. mandrillaris-infected*, untreated (vehicle only) tissues show widespread damage, particularly at edges where loss of cell density and disorder of tissue structure is apparent. Large numbers of *B. mandrillaris* trophozoites are observed intermixed with human cells and outside of the tissue (lower half of image). (C) *B. mandrillaris-infected* tissues treated with 35 μM nitroxoline simultaneously with inoculation do not show signs of tissue damage or loss of cell density and maintain distinct edges similar to uninfected tissues. Clusters of *B. mandrillaris* cysts are observed outside the boundaries of the tissue.

## Discussion

GAE caused by *Balamuthia mandrillaris* is almost always fatal (19–34). Because there is no established treatment for GAE, patients commonly receive experimental combinations of antimicrobial agents in aggressive and prolonged treatment regimens with mixed outcomes (7, 19, 23, 27, 29, 30, 34, 53–61). In the present study, we aimed to address the critical need for new treatments of GAE by identifying novel amoebicidal compounds amongst known drugs using a high-throughput screening approach. Because our goal was to identify candidate drugs with established safety and pharmacodynamic profiles, we chose to screen a library of 2,177 clinically-approved compounds. Based on the criteria of B-score and percent inhibition of *B. mandrillaris* viability, we selected 12 compounds from the primary screen for follow-up screening (Fig. 1A, Table S1B). Secondary screening eliminated all candidate compounds on the basis of low potency for inhibition of *B. mandrillaris* or high toxicity to human cells with the exception of the quinoline antibiotic nitroxoline. Nitroxoline demonstrated promising potency and selectivity for *B. mandrillaris* inhibition, which led us to focus our efforts on further investigating its novel amoebicidal activity.

We performed side-by-side experiments to directly compare the *in vitro* efficacy of nitroxoline for *B. mandrillaris* inhibition to the efficacy of pentamidine isethionate, miltefosine, and azithromycin, three drugs recommended by the CDC for treatment of GAE and commonly used in case reports (29, 30, 52–54, 56). We found that nitroxoline was the most potent inhibitor of *B. mandrillaris* trophozoite viability with an IC_50_ of 2.84 μM and IC_99_ of 7.54 μM (Fig. 3A). To estimate the selectivity of each drug, we also measured the toxicity of each compound to five different human cell lines. For pentamidine, miltefosine, and azithromycin, drug concentrations that reduced *B. mandrillaris* viability also caused significant toxicity to human cells, giving very small selectivity indices (Fig. 3B). In contrast, nitroxoline had an IC_50_ for *B. mandrillaris* inhibition that was lower than the CC_50_ for all cell lines tested and had the largest mean selectivity index.

*B. mandrillaris* and other free-living amoebae have a dormant, thick-walled cyst form which is highly resistant to several types of environmental stress including exposure to some compounds that are toxic to the trophozoite form (44, 45, 47, 49). A previous study postulates that the transition from the trophozoite form to the cyst form can be triggered by a variety of conditions including chemical stress (70), raising the possibility that treatment with amoebicidal or amoebistatic drugs may induce encystment. To investigate this possibility, we quantified the frequency of cysts observed in *B. mandrillaris* populations treated with different drug concentrations. We observed that low micromolar doses of nitroxoline and pentamidine isethionate caused an increase in the total number and proportion of cysts in *B. mandrillaris* populations (Fig. 4A and B). In contrast, miltefosine and azithromycin were not observed to induce encystment at any concentration (Fig. 4C and D). These data support the possibility that *B. mandrillaris* encystment occurs as a response to some but not all compounds that are toxic to trophozoites. We suggest that compounds which promote encystment in addition to killing trophozoites may provide additional benefit in the context of infection by slowing or halting the rapid tissue destruction by trophozoites.

Because it is common for both cysts and trophozoites to be found in *B. mandrillaris* infected tissue, it is important to understand the efficacy of amoebicidal drugs against both stages (71). To address this question, we performed side-by-side dose-response experiments with homogenous populations of cysts or trophozoites treated with nitroxoline, pentamidine isethionate, miltefosine, and azithromycin. We found that cysts were less sensitive than trophozoites to all four drugs (Fig. 4E). The difference in sensitivity was marginal for miltefosine and greatest for nitroxoline, which had an IC_50_ for cyst inhibition 6-fold higher than the IC50 for trophozoite inhibition. Despite being less potent against the cyst form, nitroxoline still exhibited the lowest IC_50_ for cyst inhibition of all drugs tested at 15.48 μM.

Viability measurements of drug treated *B. mandrillaris* cultures reflect a complex population phenotype that includes varying degrees of both death and encystment. These measurements are not sufficient to predict the rate at which populations will recrudesce following treatment, which is an important factor in evaluating the overall efficacy of different treatments. We chose to address this aspect of treatment efficacy by performing recovery assays in which *B. mandrillaris* populations were exposed to various drug treatments, removed from drug, and then co-cultured with monolayers of human cells until the host cells were fully consumed. As predicted, increasing drug doses caused greater delays to *B. mandrillaris* recrudescence and host cell destruction (Table 1). Surprisingly, even low micromolar doses of nitroxoline near the IC_50_ for trophozoite inhibition delayed *B. mandrillaris-mediated* host cell destruction by 2-3 weeks. This assay also served as a sensitive method to detect very low numbers of surviving amoebae due to the large population expansion occurring over the 28-day experiment duration. Importantly, this sensitive assay allowed us to infer that *B. mandrillaris* populations had been completely eliminated by drug treatment when we observed no signs of amoeba population recovery after 28 days of co-culture with host cells.

**Table 1:** Rate of recrudescence and host cell destruction by *B. mandrillaris* following drug treatment. Cultures of *B. mandrillaris* were exposed to the indicated drug concentrations for 72 hours, removed from drug, and transferred to monolayers of human cells where the time until complete monolayer clearance was measured. Mean host cell clearance time is reported as the average of three replicates ± the standard error. Low micromolar doses of nitroxoline delay *B. mandrillaris* recrudescence by two to three weeks compared to untreated controls. Doses of 28 μM nitroxoline and 112 μM miltefosine completely prevented recovery of *B. mandrillaris* and damage to host cells throughout the 28-day duration of the experiment.

| Treatment | Mean days for host cell clearance ± SE |
| --- | --- |
| Vehicle only |  |
| 0.28% DMSO v/v | 1.33 ± 0.33 |
| 1.12% Water v/v | 1.67 ± 0.67 |
| Nitroxoline (in DMSO) |  |
| 3.5 μM | 15.33 ± 0.67 |
| 7 μM | 19 ± 0.58 |
| 14 μM | 22 ± 2.08 |
| 28 μM | NRT<sup>a</sup> |
| Pentamidine isetionate (in water) |  |
| 3.5 μM | 1.33 ± 0.33 |
| 7 μM | 7 ± 1.15 |
| 14 μM | 13 ± 2.51 |
| 28 μM | 15.33 ± 2.56 |
| 56 μM | 16 ± 2 |
| Miltefosine (in water) |  |
| 28 μM | 1.5 ± 0.33 |
| 56 μM | 3.5 ± 2.64 |
| 84 μM | >28<sup>b</sup> |
| 112 μM | NRT<sup>a</sup> |

<sup>a</sup> No recovery of trophozoites or host cell destruction observed through 28 days
<sup>b</sup> Incomplete recovery of trophozoites and destruction of host cells on day 28

Using this method, we determined that doses of 28 μM nitroxoline and 112 μM miltefosine completely eliminated *B. mandrillaris* populations. These data are consistent with viability inhibition experiments in indicating that nitroxoline is the most potent inhibitor of *B. mandrillaris* tested. The promising results of this experiment suggest that nitroxoline may be able to fully eliminate *B. mandrillaris* infection if high enough concentrations are reached and is likely to cause substantial delays to host tissue damage even at lower concentrations. This is supported by our findings that nitroxoline prevents *B. mandrillaris* activity and tissue destruction in a primary human brain tissue model (Fig. 5, Movie S1 and S2). Together, these findings suggest that nitroxoline substantially impedes host tissue destruction by *B. mandrillaris in vitro*. Given the rapid progression of pathogenesis that is characteristic of GAE, any impediment to tissue destruction could significantly improve patient prognosis.

The pharmacokinetic properties of nitroxoline suggest further promise for its therapeutic value as an inhibitor of *B. mandrilliaris*. Nitroxoline has been safely used for over 50 years in the treatment of urinary tract infections with minimal adverse effects reported (72). Nitroxoline is available in oral and intravenous administration and is typically dosed at 600-800 mg/day for adults, resulting in maximal plasma concentrations (C_max_) up to approximately 30 μM (5.6 mg/L) (62–64), which is 10-fold higher than the IC_50_ for *B. mandrillaris* trophozoites *in vitro* (Fig. 3A). Although the extent to which nitroxoline crosses the blood brain barrier is unknown, a recent study shows that systemically delivered nitroxoline exhibits efficacy against gliomas in mice, implying that efficacious concentrations reached the brain in this model (73). In addition, we previously noted that *B. mandrillaris* frequently causes a necrotizing vasculitis in the CNS with extensive BBB breakdown (3, 36, 56). As a result, the bioavailability of nitroxoline in the CNS will almost certainly be significantly increased in patients actively suffering from GAE. Furthermore, given the extreme severity of GAE, intrathecal drug delivery can be performed to maximize drug concentrations reaching the brain (2, 74). While many variables may affect the *in vivo* efficacy of nitroxoline as well as the concentrations that can be achieved in relevant compartments, the literature suggests that the *in vitro* efficacious concentrations we demonstrate are well within a physiologically relevant range.

The data presented in this study strongly indicate that nitroxoline warrants further investigation as a potential treatment for *B. mandrillaris* infections. As an approved compound with an established safety profile (72), nitroxoline may be rapidly deployed as an experimental treatment in dire cases of GAE. In particular, since the current standard of care generally consists of experimental combinations of several antimicrobial agents, adding nitroxoline to these regimens may be a reasonable step in the effort to improve patient prognosis. Although our SAR experiments did not identify any analog that matched the potency of nitroxoline, medicinal chemistry optimization may still be beneficial for a better understanding of possible mechanisms of action and efforts to improve drug specificity. The similarity of *B. mandrillaris* to other free-living amoebae such as *Acanthamoeba spp*. and *Naegleria fowleri* raises the intriguing possibility that nitroxoline or related compounds may also have activity against these pathogens.

## Author contributions

Study design and conception: MTL, CVW, JLD Data acquisition: MTL, CVW, HR, MSM Data analysis and interpretation: MTL, CVW, HR, WW, JS, HA, CW, MRA, JLD Drafting of manuscript: MTL, CVW, JS, and JLD Critical revision of manuscript: All authors

## Acknowledgements

We thank Jiri Gut for critical advice regarding the axenic culture of *Balamuthia* mandrillaris. We are grateful to Galina Schmunk, Tom Nowakowski, and Arnold Kriegstein for their assistance with human tissue experiments. We thank Michael Wilson for helpful discussion and critical reading of the manuscript.

---

**Table S1:** Tabulation of raw data from primary and secondary drug screens. (A) A primary screen of 2,177 clinically-approved compounds measured the decrease in *B. mandrillaris* trophozoite viability resulting from 72 hours of exposure to 20 μM of each compound. Viability for test and control wells was measured using the CellTiter Glo luminescence assay. Raw luminescence intensity values for test wells are reported as well as the average luminescence intensity values for positive and negative controls used for comparison. The B-score and percent inhibition shown for each compound are calculated using the raw luminescence intensity values. (B) List of compounds meeting loosely applied criteria of B-score of approximately 5 or greater and percent inhibition of approximately 40 or greater. The class and delivery method for each compound is annotated. Compounds that are classified as veterinary or are available only for external/topical administration are highlighted in yellow. Nitroxoline, the top hit compound based on B-score and percent inhibition, is highlighted in red. (C) Secondary screening of hit compounds to determine potency against *B. mandrillaris* trophozoites and toxicity to HFF1 and H4 human cell lines. All IC_50_ values were approximated from 8-point dose-response curves. IC_50_ values highlighted in red are outside of a desirable range for a hit compound as they indicate either low potency against *B. mandrillaris* (IC_50_ > 30 μM) or high toxicity to human cells (IC_50_ < 1 μM). Nitroxoline and pentamidine isethionate were the only compounds that inhibited *B. mandrillaris* trophozoite viability at concentrations similar to or lower than concentrations that were toxic to human cells.

**Table S2:**
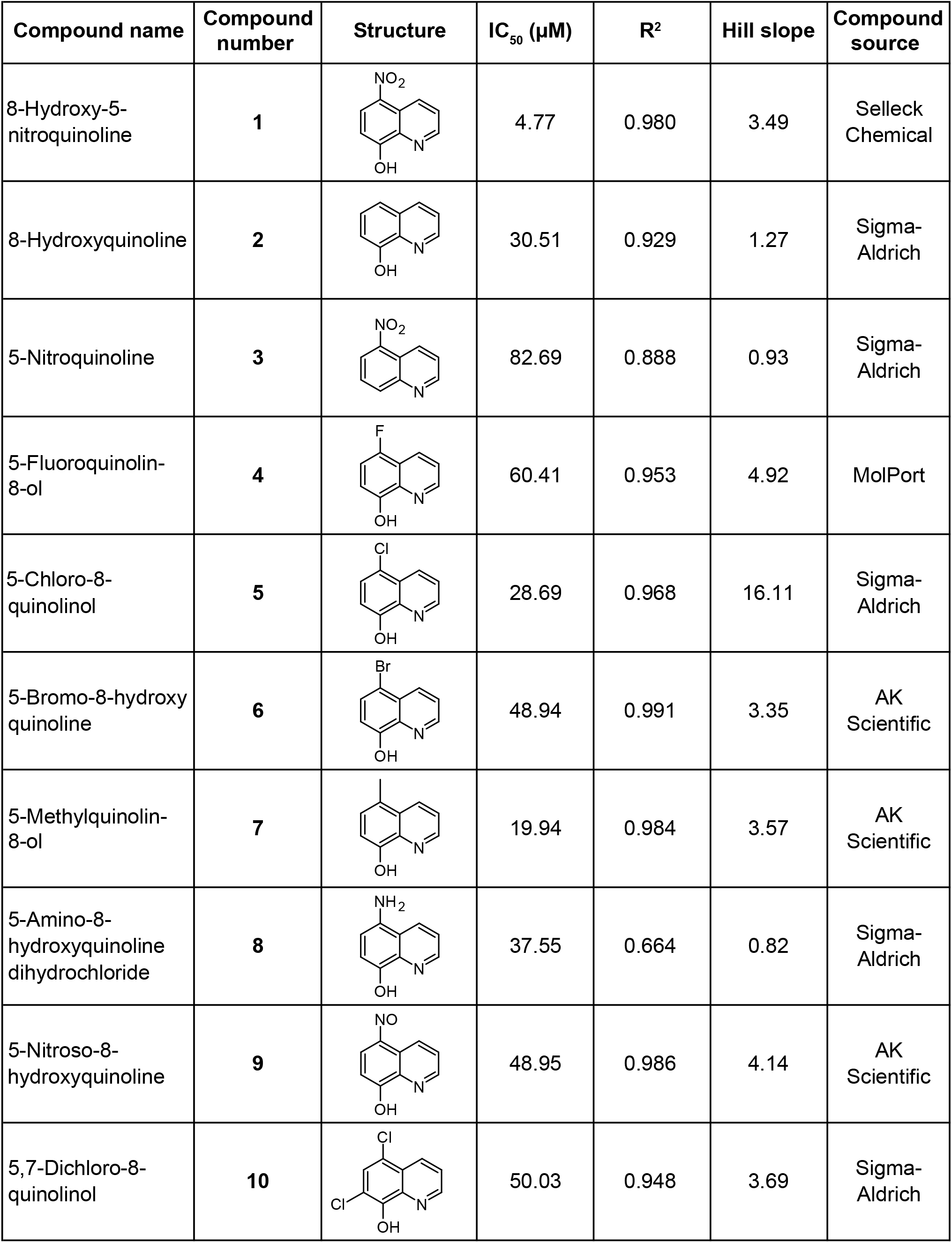
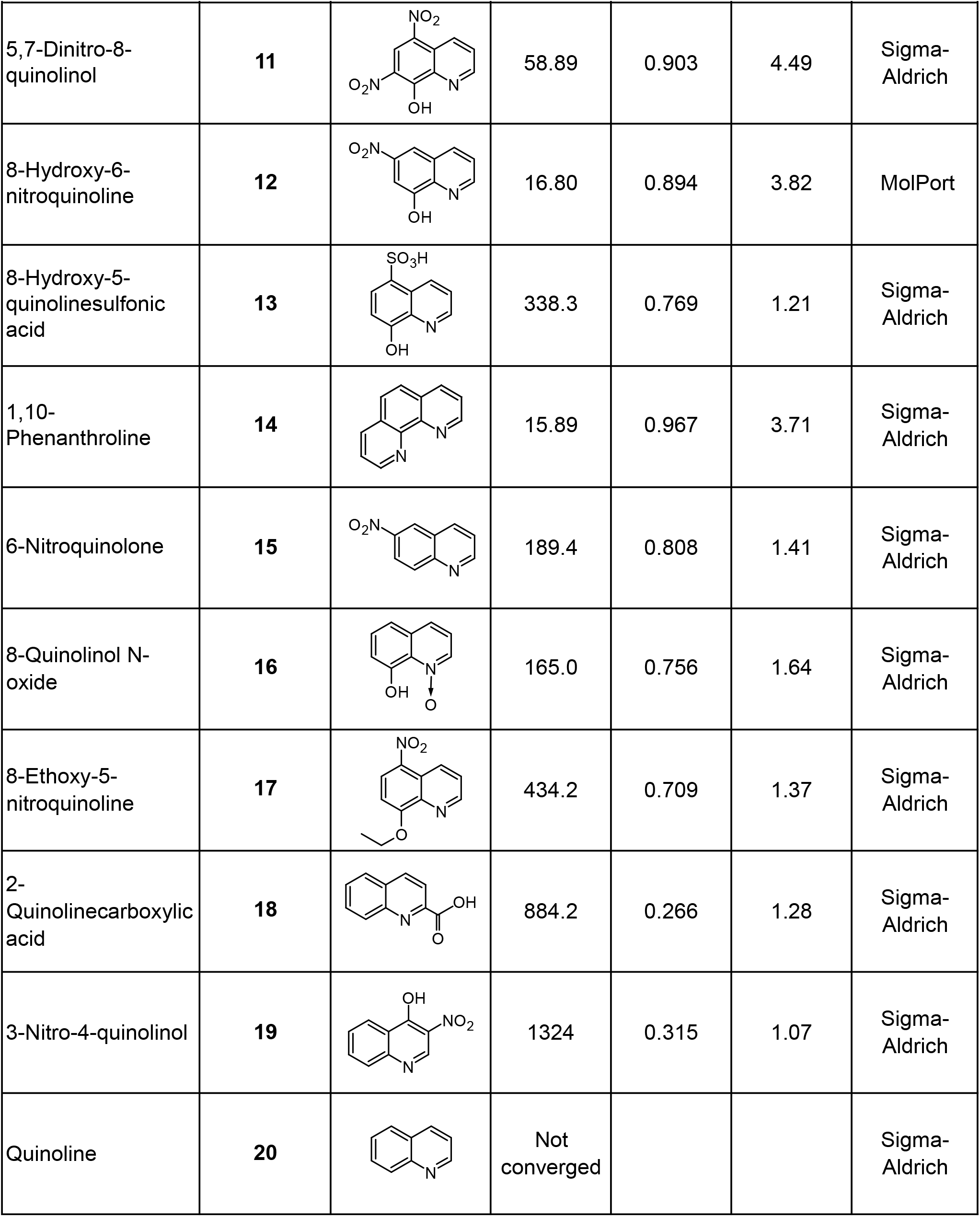
Structures, IC_50_ values, and dose-response curve fitting parameters for nitroxoline and all nitroxoline analogs tested for activity against *B. mandrillaris*. IC_50_, R^2^, and Hill slope values were calculated based on sigmoidal curves fit to 8-point dose-response experiments performed in triplicate. Compounds 2, 4-12, and 14 were the most potent inhibitors of *B. mandrillaris*, most likely due to the presence of an 8-position hydroxyl group or bidentate ligand on the quinoline ring, which are predicted to be necessary for metal binding activity.

| Compound name | Compound number | Structure | IC<sub>50</sub> (μM) | R<sup>2</sup> | Hill slope | Compound source |
| --- | --- | --- | --- | --- | --- | --- |
| 8-Hydroxy-5-nitroquinoline | **1** |  | 4.77 | 0.980 | 3.49 | Selleck Chemical |
| 8-Hydroxyquinoline | **2** |  | 30.51 | 0.929 | 1.27 | Sigma-Aldrich |
| 5-Nitroquinoline | **3** |  | 82.69 | 0.888 | 0.93 | Sigma-Aldrich |
| 5-Fluoroquinolin-8-ol | **4** |  | 60.41 | 0.953 | 4.92 | MolPort |
| 5-Chloro-8-quinolinol | **5** |  | 28.69 | 0.968 | 16.11 | Sigma-Aldrich |
| 5-Bromo-8-hydroxyquinoline | **6** |  | 48.94 | 0.991 | 3.35 | AK Scientific |
| 5-Methylquinolin-8-ol | **7** |  | 19.94 | 0.984 | 3.57 | AK Scientific |
| 5-Amino-8-hydroxyquinoline dihydrochloride | **8** |  | 37.55 | 0.664 | 0.82 | Sigma-Aldrich |
| 5-Nitroso-8-hydroxyquinoline | **9** |  | 48.95 | 0.986 | 4.14 | AK Scientific |
| 5,7-Dichloro-8-quinolinol | **10** |  | 50.03 | 0.948 | 3.69 | Sigma-Aldrich |
| 5,7-Dinitro-8-quinolinol | 11 |  | 58.89 | 0.903 | 4.49 | Sigma-Aldrich |
| 8-Hydroxy-6-nitroquinoline | 12 |  | 16.80 | 0.894 | 3.82 | MolPort |
| 8-Hydroxy-5-quinolinesulfonic acid | 13 |  | 338.3 | 0.769 | 1.21 | Sigma-Aldrich |
| 1,10-Phenanthroline | 14 |  | 15.89 | 0.967 | 3.71 | Sigma-Aldrich |
| 6-Nitroquinolone | 15 |  | 189.4 | 0.808 | 1.41 | Sigma-Aldrich |
| 8-Quinolinol N-oxide | 16 |  | 165.0 | 0.756 | 1.64 | Sigma-Aldrich |
| 8-Ethoxy-5-nitroquinoline | 17 |  | 434.2 | 0.709 | 1.37 | Sigma-Aldrich |
| 2-Quinolinecarboxylic acid | 18 |  | 884.2 | 0.266 | 1.28 | Sigma-Aldrich |
| 3-Nitro-4-quinolinol | 19 |  | 1324 | 0.315 | 1.07 | Sigma-Aldrich |
| Quinoline | 20 |  | Not converged |  |  | Sigma-Aldrich |

**Figure S1:**
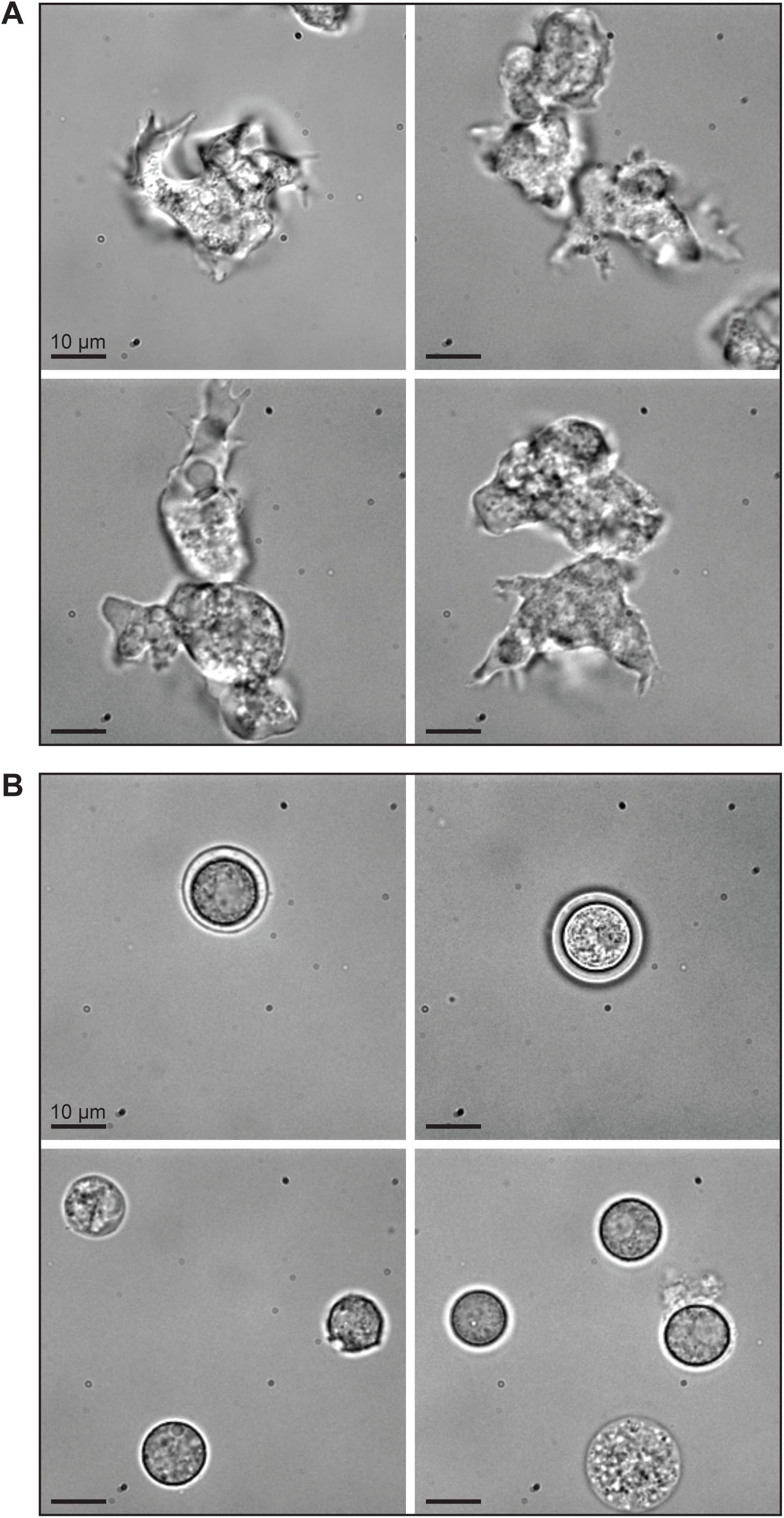
Example brightfield images of *B. mandrillaris* trophozoites and cysts. (A) Example images show *B. mandrillaris* trophozoites in log-phase growth. Trophozoites are pleomorphic and can be elongated or generally rounded, often with highly branched pseudopodia. (B) Example images show *B. mandrillaris* cysts induced by galactose exposure. Cysts are spherical, generally smaller in diameter than trophozoites, and can have visibly distinct layers. Some cysts show signs of vacuolization that may be indicative of cell death.

**Figure S2:**
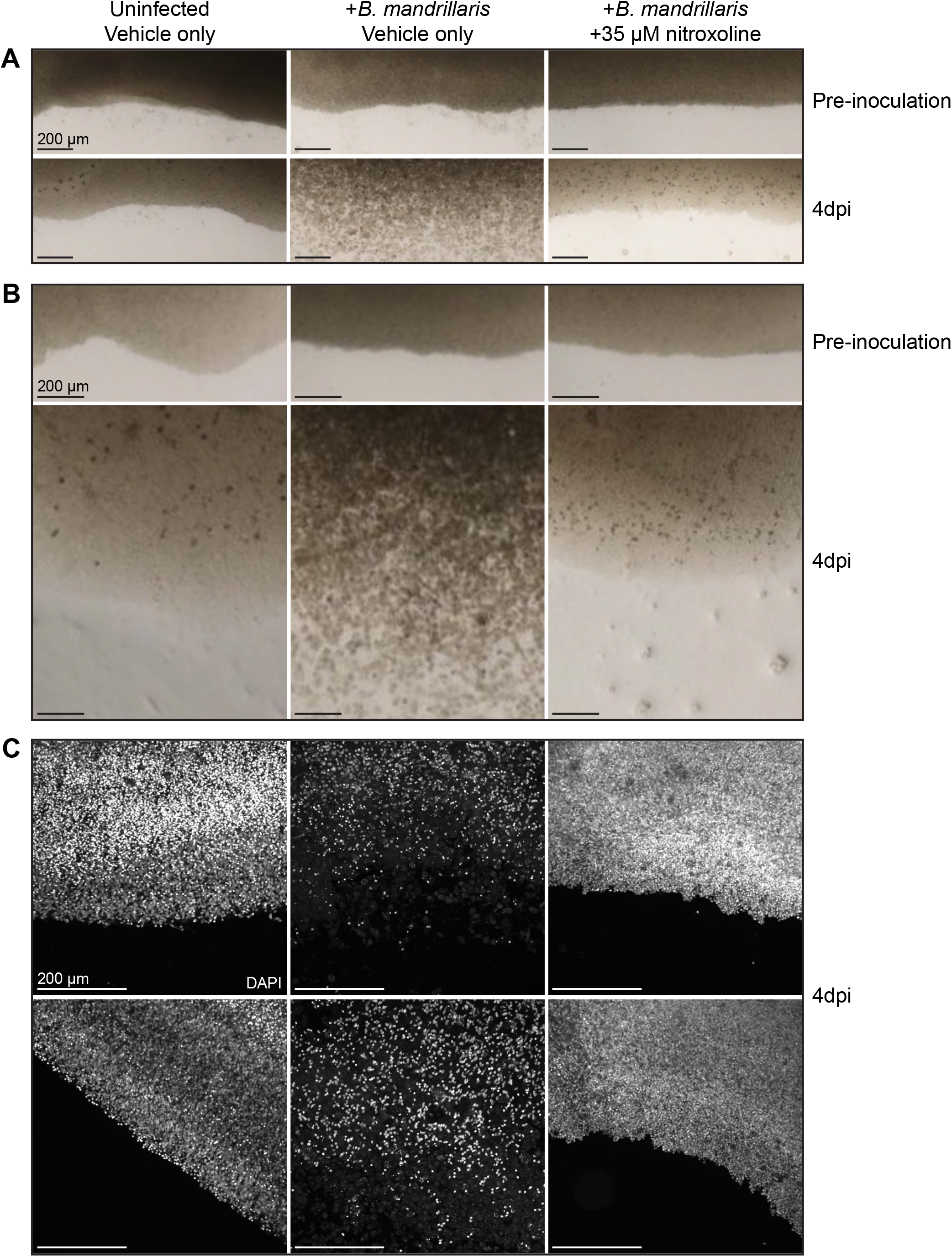
(A and B) Brightfield images representative of two biological replicates for human brain tissue explants before and after exposure to the indicated conditions. Nitroxoline or vehicle was added simultaneously with *B. mandrillaris* trophozoites and removed after 20 hours. Four days after exposure to *B. mandrillaris*, untreated tissues show widespread damage and loss of cell density while nitroxoline treated tissues remain intact and appear similar to uninfected tissues. Large numbers of *B. mandrillaris* trophozoites can be seen at the edges of untreated tissues, while only clusters of cysts are observed in nitroxoline-treated tissues. (C) Images representative of two biological replicates of human brain tissue explants fixed and stained with DAPI four days after exposure to the indicated conditions (two images per condition). The number of host cell nuclei is dramatically reduced in untreated, *B. mandrillaris-infected* tissues compared to uninfected tissues, whereas *B. mandrillaris-infected* tissues treated with nitroxoline show no apparent loss of nuclei.

**Movie S1:** (A) Representative movie showing the edge of an untreated (vehicle only), uninfected primary brain tissue section 4 days after the start of culture. No significant movement of cells is observed and a clearly defined edge marks the border of the tissue. (B and C) Representative movies showing edges of two different untreated (vehicle only), *B. mandrillaris-infected* primary brain tissue sections 4 days after inoculation of amoebae. Highly motile *B. mandrillaris* trophozoites are observed throughout the field of view and the tissue edges are not discernable due to loss of host cell density. (D and E) Representative movies showing edges of two different nitroxoline treated, *B. mandrillaris-infected* primary brain tissue sections 4 days after inoculation of amoebae. Tissues were treated with 35 μM nitroxoline simultaneously with amoebae inoculation. Non-motile *B. mandrillaris* cysts are observed individually primarily at the edge of the tissue (D) and in clusters outside the tissue (E). The tissue edges are intact and do not show signs of host cell destruction by *B. mandrillaris*.

